# Small molecule NS11021 promotes BK channel activation by increasing inner pore hydration

**DOI:** 10.1101/2024.06.03.597166

**Authors:** Erik B. Nordquist, Zhiguang Jia, Jianhan Chen

## Abstract

The Ca^2+^ and voltage-gated big potassium (BK) channels are implicated in various diseases including heart disease, asthma, epilepsy and cancer, but remains an elusive drug target. A class of negatively charged activators (NCAs) have been demonstrated to promote the activation of several potassium channels including BK channels by binding to the hydrophobic inner pore; yet the underlying molecular mechanism of action remains poorly understood. In this work, we analyze the binding mode and potential activation mechanism of a specific NCA named NS11021 using atomistic simulations. The results show that NS11021 binding to the pore in deactivated BK channels is nonspecific and dynamic. The binding free energy of -8.3±0.7 kcal/mol (*KD* = 0.3-3.1 μM) calculated using umbrella sampling agrees quantitatively with the experimental EC50 range of 0.4-2.1 μM. The bound NS11021 remains dynamic and is distal from the filter to significantly impact its conformation. Instead, NS11021 binding significantly enhances the pore hydration due to the charged tetrazol-phenyl group, thereby promoting the opening of the hydrophobic gate. We further show that the free energy barrier to K^+^ permeation is reduced by ∼3 kcal/mol regardless of the binding pose, which could explain the ∼62-fold increase in the intrinsic opening of BK channels measured experimentally. Taken together, these results support that the molecular mechanism of NS11021 derives from increasing the hydration level of the conformationally closed pore, which does not depend on specific binding and likely explains the ability of NCAs to activate multiple K^+^ channels.

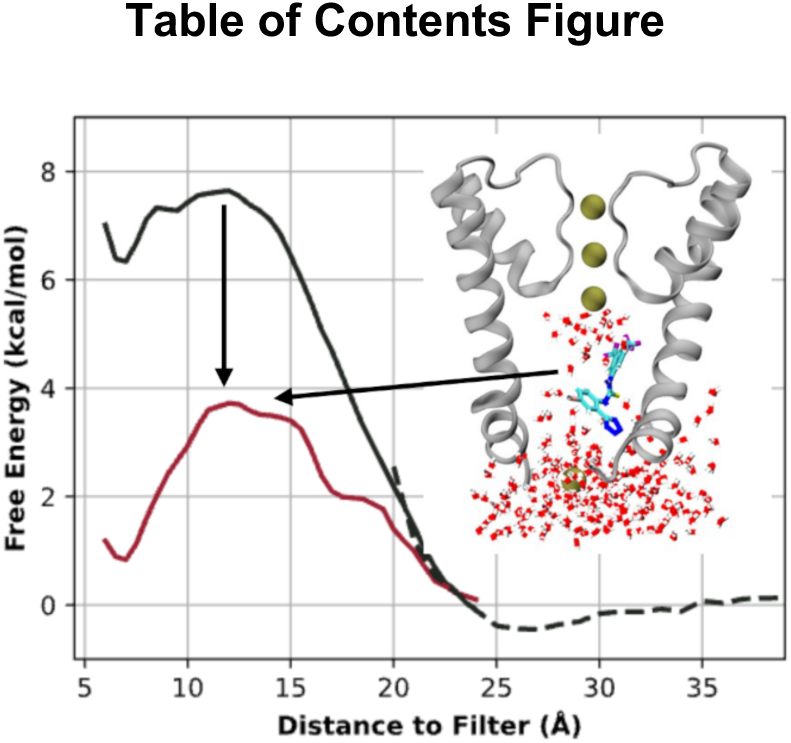

## Introduction

Ion channels are an important class of drug targets and are the targets for ∼20% of drugs with FDA approval.^1^ The Ca^2+^ and voltage-gated K^+^ channel (Figure 1A), also referred to as KCa1.1, MaxiK, or big potassium (BK) channel,^2–4^ has been extensively studied for its role in the nervous system,^5^ smooth muscle contraction,^6,7^ and hearing.^8^ It has been implicated in numerous diseases, including stroke,^9–11^ ischemia-reperfusion injury,^12,13^ asthma,^14–16^ epilepsy,^17–20^ urinary bladder disorder,^21–23^ rheumatoid arthritis^24,25^ and cancers of numerous types.^26–28^ Several BK channel activators are on the market for treating breast cancer,^29^ epilepsy,^30,31^ and blood flow,^32^ and Andolast has been approved in Italy as asthma medication.^33^ Despite these advances, the BK channel remains a challenging drug target and there are no compounds with FDA approval that directly target the BK channel. This deficit is not for a lack of tool compounds with known activity for the BK channel.^34–36^ Rather, it is due to challenges such as designing molecules specific for a tissue-localizing β or γ subunit and achieving sufficient potency yet minimal off-target effects.^37,38^ Another important challenge lies in limited understanding of the molecular mechanisms of action of known compounds.

**Figure 1:**
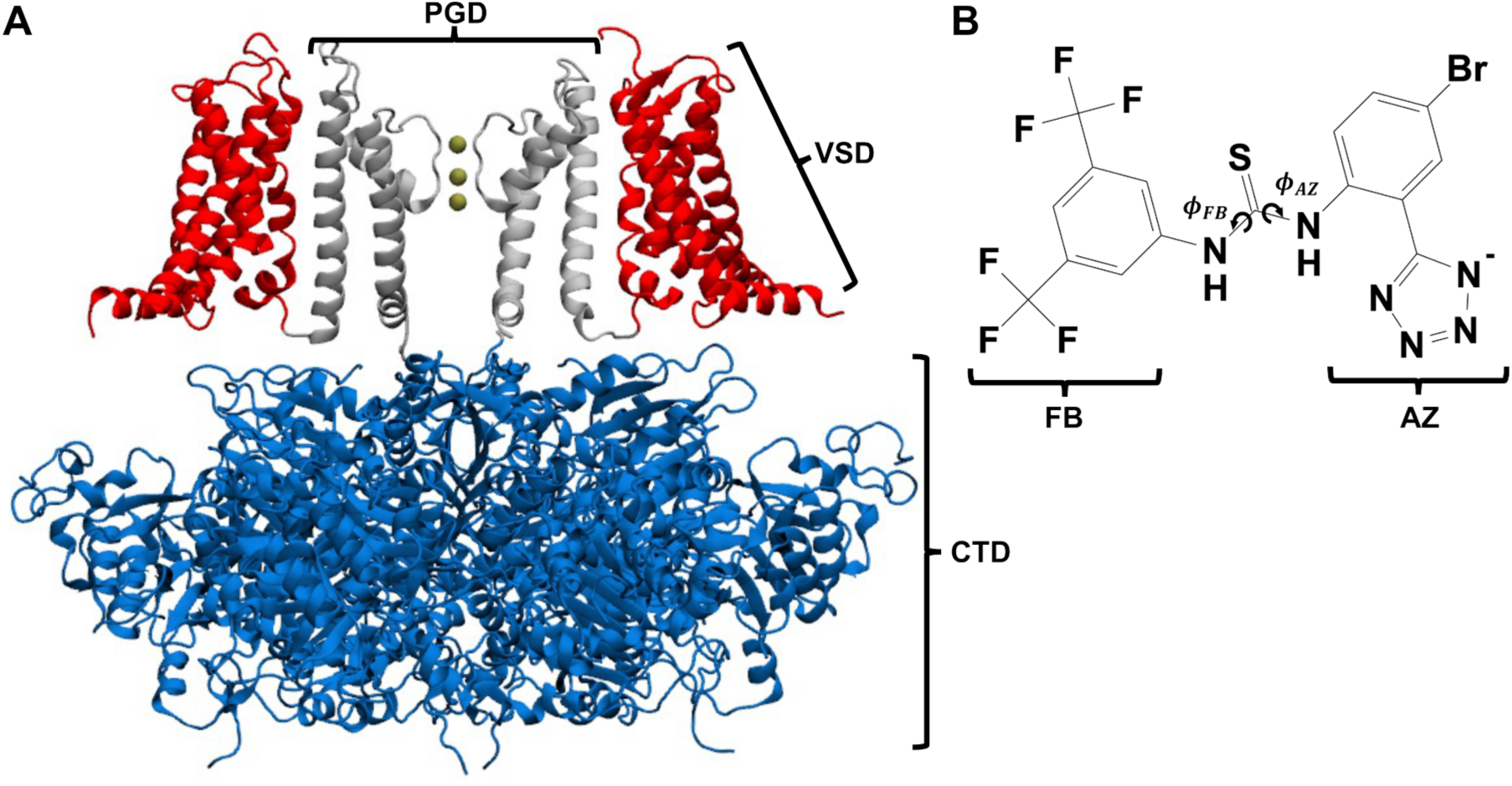
Structures of the BK channel and activator NS11021. A) Human BK channel α subunit in the Ca^2+^-free, presumably deactivated state (PDB: 6v3g^39^). For clarity, only two of the four VSD (red) and PGD (silver) are rendered. The CTD is blue and the K^+^ in the selectivity filter are gold. **B)** Chemical structure of NS11021, with tetrazol-phenyl (AZ) and trifluoromethyl- phenyl (FB) groups labelled. Partial charges of the AZ group are shown in Figure S1.

The BK channel α subunit is a homotetramer, composed of a transmembrane (TM) region which houses two domains, and a cytosolic tail domain (CTD) primarily responsible for sensing Ca^2+^ (Figure 1A). The TM region includes a voltage sensing domain (VSD), which consists of five TM helices S0-S4, and pore gating domain (PGD), which consists of the two innermost TM helices S5-S6 and the K^+^ selectivity filter (Figure 1A). The smooth muscle relaxant NS11021 (Figure 1B),^21,40,41^ is a BK channel activator known to act on the pore.^42^ It belongs to a class of negatively-charged activators (NCA) of K2P, hERG and BK channels.^43^ These NCAs share a common pharmacophore that contains a negatively charged group attached to aromatic and hydrophobic moieties. X-ray crystallography together with cysteine-scanning mutagenesis and molecular docking revealed that the NCA BL-1249 binds to the TREK-2 K2P channel in the pore with the negatively-charged group pointed towards the selectivity filter.^43^ The binding location of BL-1249 was inferred from the location of a Br atom in BL-1249, as no other electron density corresponding to BL-1249 was observed.^43^ Patch-clamp electrophysiology experiments revealed that BL-1249 increases the single-channel conductance of TREK-2,^43^ putatively by inducing a change in selectivity filter conformation.^44^ Atomistic simulations further suggested that the increase in conductance results from the stabilization of K^+^ binding beneath the filter (in the “S6” site).^43^ Similar positioning of K^+^ ions was previously shown via atomistic simulation to alter the conformation of the selectivity filter, thereby opening the filter gate of K2P channels.^44^ It was proposed that the NCA NS11021 acts on the BK channel via a similar mechanism.^43^

There are several key experimental observations which inform how NS11021 acts on the BK channel. Rockman, *et al.* showed that NS11021 activity had little Ca^2+^-dependence and that truncation of the CTD did not significantly alter its activity, suggesting that NS11021 does not interact with the CTD.^45^ Furthermore, NS11021 activity has minimal voltage-dependence, suggesting that it has minimal impact on VSD activation or VSD-PGD coupling.^45^ By fitting many voltage-dependent dose-response curves with the Hill equation, they showed that the molecule binds in a 1:1 molar ratio with the channel. Together, these electrophysiological analyses strongly support the notion that the molecule binds in the central pore and acts on the PGD itself, consistent with the observation from X-ray crystallography.^43^ The authors further performed limiting slope experiments, where the voltage-dependence of the channel’s intrinsic open probability *PO* was measured down to -200 mV in 0 Ca^2+^, a regime in which the VSD is resting. The results reveal that NS11021 increases the *P_O_* by 62-fold at 30 µM. Therefore, the primary effect of NS11021 is to shift the pore open-close equilibrium towards the open state, by increasing the average open state dwell time while substantially decreasing the average dwell time of closed states.^45^ However, NS11021 does not significantly the increase single channel conductance,^45^ which suggests that the activity of NS11021 in BK is not likely through the putative filter-gate previously proposed.^43^

In this study we use atomistic molecular dynamics (MD) simulations to examine the molecular detail and free energy of NS11021 binding to the inner pore of the BK channel in the deactivated state. The results show that the NS11021 binds nonspecifically to the hydrophobic surface of the pore and remains highly dynamic inside the pore, which could explain a lack of resolvable density for NCA BL-1249 in TREK2 discussed above.^43^ The calculated free energy of binding of ∼ -8 kcal/mol is consistent with published experimental EC50 measurements.^42,45^ Importantly, the presence of NS11021 in the pore significantly increase the level of hydration. This effect is significant because it has been proposed that BK channels likely followed the hydrophobic gating mechanism, where the pore undergoes a spontaneous dewetting transition to deactivate the channel.^46–48^ Increasing the pore hydration will stabilize the hydrated open state, as suggested by electrophysiological analysis.^45^ Furthermore, analysis of K^+^ permeation free energy profiles with NS11021 in three representative binding conformations show that the free energy barrier is reduced by 3-5 kcal/mol compared to the empty pore. This is in good agreement with the experimentally-derived 62-fold shift in the intrinsic *PO* induced by NS11021,^45^ which corresponds to a 2.5 kcal/mol decrease in the K^+^ permeation barrier. Taken together, these results strongly support that NS11021 promotes BK activation by lowering the barrier to pore hydration instead of direct modulation of the putative filter gate.

## Methods

### Parameterization of small molecule NS11021

Molecular model parameters for the NS11021 ligand were assigned by identifying similar atom, bond, angle, dihedral types as well as charges from similar small molecules within the CHARMM General Force Field (CGenFF).^49,50^ The only substitution group absent in the CGenFF force field is the tetrazol-phenyl group. The structure of this group was optimized at the MP2/6-31G* level using the program GAMESS,^51–53^ and the atomic charges were then calculated using the Merz-Singh-Kollman scheme^54^ as implemented in the Multiwfn program^55^ (Figure S1). The CGenFF parameters released in July 2022 included a methylthiourea torsional profile with barriers of 5 kcal/mol. We simulated a single NS11021 molecule in CHARMM TIP3P water for 100 ns using the July 2022 thiourea dihedral parameters and observed many reversible transitions. All four *syn-* and *anti-* combinations (Figure S2) were accessible with stabilities differing by ∼1-3 kcal/mol (Figure S3), in qualitative agreement with solution NMR data of diaryl-thiourea compounds.^56^ The topology and parameters used for the whole system, including the NS11021 ligand, are available on GitHub at https://github.com/enordquist/BK_NS11021_SI/.

### Atomistic simulations of BK channels

The BK channel structure was derived from the apo-structure 6v3g.^39^ We used the same system configuration as previously described, including initial minimization, seven steps of NVT equilibration while iteratively lowering restraints on the system’s components, and additional relaxation simulations to allow a spurious π-helix bulge on the pore-lining S6 helix to spontaneously refold back to a continuous α-helix.^48^ We performed all simulations using only the TM region and C-linkers to mimic the Core-MT construct,^57,58^ which retains voltage gating but has no Ca^2+^ dependence. All Cα atoms of the protein were harmonically restrained with a force constant of 0.2 kcal/mol/Å^2^ during free energy calculations. The simulations were performed using the MPI-enabled GROMACS version 2019^59,60^ and PLUMED version 2.7.^61,62^ The initial simulation box with a POPC lipid bilayer, TIP3P water and 150 mM KCl plus neutralizing ions was generated using the CHARMM-GUI webserver.^63,64^ The final simulation boxes were approximately of dimension 160 × 160 × 100 Å, and contained ∼270,000 atoms. We observed frequent reversible lipid entry through the fenestration window between S6 helices, as observed consistently in previous simulations and multiple cryo-EM maps.^39,46–48,65^ Occasionally, some lipid tails may remain in the pore for tens of ns. To avoid sampling these infrequent events during umbrella sampling free energy calculations (see below), a linear restraint of the form *k N_lipid_* with k = 40 kcal/mol was imposed to prevent the lipids from fully entering the pore using the PLUMED collective variable (CV) COORDINATION, as previously described.^48^

The systems were described using the CHARMM36m protein^66,67^ and CHARMM36 lipid^68^ force fields. Lengths of all hydrogen-containing bonds were constrained with the LINCS algorithm,^69,70^ the MD timestep was 2 fs, Electrostatics were treated with the particle mesh Ewald algorithm^71^ using a 12 Å distance cutoff, and Van der Waals forces were smoothly switched off from 10 to 12 Å. The Nosé-Hoover thermostat^72,73^ was used with reference temperature of 303.15 K and coupling constant *τ_T_* of 1 ps^−1^. Constant pressure was imposed using the Parrinello-Rahman^74,75^ semi-isotropic barostat in *x* and *y* directions using reference pressure of 1 bar, compressibility of 4.5 × 10^−5^ bar^−1^, and pressure coupling constant *τ_P_* of 5 ps^−1^.

### Umbrella sampling of NS11021 binding to the pore

Pilot simulations show that NS11021 had sufficient space in the pore to sample a wide range of orientations and configurations, even though the exchange appeared relatively slow compared to umbrella sampling timescales (up to 100 ns per window). Accordingly, we performed umbrella sampling simulations in two parallel sets, initiating each set of windows with either the (3,5)- FB or AZ group facing up (towards the pore and filter) (Figure 1B). Sampling from both sets of windows will be combined to derive the overall free energy surface. In each umbrella sampling window, harmonic distance restraint potential was imposed on the z-distance between the centers-of-mass (CoM) of NS11021 and the selectivity filter with a force constant of 0.5 kcal/mol/Å^2^. The CoM of the selectivity filter is determined using the backbone heavy atoms of residues 286, 287 and 288, which effectively corresponds to the S4 K^+^ binding site.^76^ To generate the initial configurations of all windows, two 8-ns steered MD simulations were performed with either the FB or AZ group pointing up towards the filter. Both sets of umbrella sampling simulations consist of five windows centered at z-distance to filter of 12, 14, 16, 18, and 22 Å, respectively. During umbrella sampling simulations, NS11201 could rarely exchange between “AZ-up” and “FB- up” orientations (Figure S4, Figure S5, and Figure S6). This adds to the sampling challenge, and thus requires long sampling time of 100 ns per window to achieve sufficient convergence in the final free energy surfaces. In addition, four windows centered at 24, 26, 29 and 32 Å were added to improve sampling in the bulk region outside the pore (Figure 2B). Finally, to improve convergence around the narrowest region of the pore where sampling of NS11021 configuration is slow, four more windows were added to the 12-16 Å region as follows: two windows centered at 14 and 16 Å, respectively with initial FB-up orientations, one window centered at 16 Å with the AZ and FB groups at the same level, and one window centered at 12 Å in an AZ-up orientation with Nwater harmonically restrained at 15 with a force constant of 0.2 kcal/mol/Nwater^2^. All windows lasted 100 ns and snapshots were saved every 5 ps. We discarded the initial 15 ns of each window and used the remaining samples to construct potentials of mean force (PMFs) using the weighted histogram analysis method (WHAM).^77^ All initial snapshots as well as representative GROMACS MD parameter (mdp) and PLUMED run files are available at https://github.com/enordquist/BK_NS11021_SI/.

**Figure 2:**
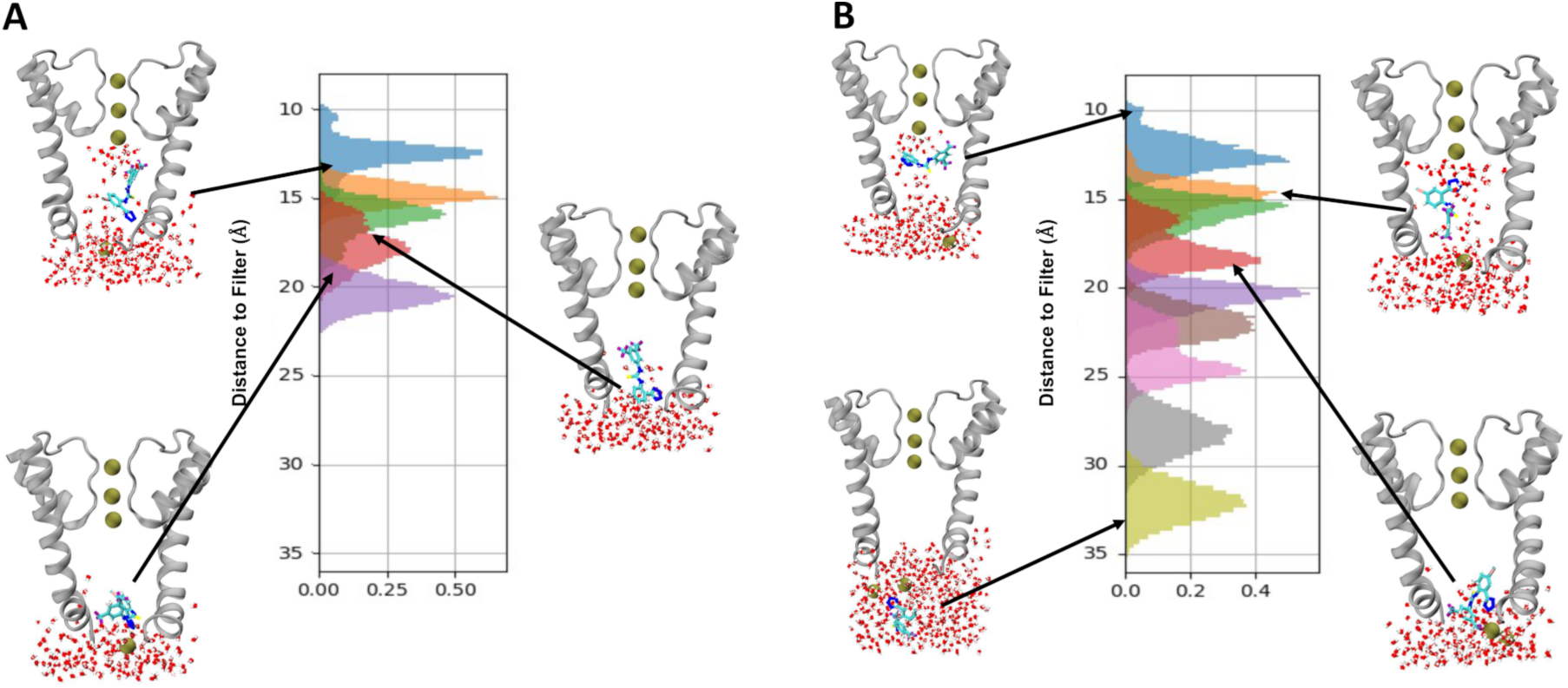
Umbrella sampling of NS11021 binding to the BK pore. Representative snapshots and z-distance to the filter histograms for the set of windows with NS11021 in the **A)** FB-up and **B)** AZ-up orientations. Panel **B)** also shows histograms and a representative snapshot from the four bulk sampling windows (see Methods). Representative snapshots depict two opposing pore-lining S6 helices and the selectivity filter (residues 273–330) in silver. Potassium ions are drawn as gold spheres, and NS11021 and water molecules are drawn as bonds, where oxygen atoms are drawn in red, hydrogen in white, carbon in cyan, nitrogen in blue, fluorine in purple, and bromine in pink. Note that the pore is largely dehydrated without NS11021.

### Calculation of K^+^ permeation free energy

To compute the K^+^ permeation PMF without NS11021, we used eight umbrella sampling windows along the K^+^ z-distance to the filter CoM as defined above from 22 Å to 8 Å spaced in 2 Å increments. The initial configurations were generated using a 2-ns steered MD simulation. In each window, the selected K^+^ was harmonically restrained using a force constant of 1.0 kcal/mol/Å^2^. All windows were run for 20 ns, and the first 1 ns was discarded in WHAM analysis. To calculate the K^+^ permeation PMF with NS11021 bound in the pore, we selected representative conformations of three distinct bindings states and harmonically restrained the CoM of NS11021 along the pore’s z-axis to the initial position with a force constant of 0.5 kcal/mol/Å^2^. This restraint allows the molecule to respond to K^+^ but prevents NS11021 from moving too far from the initial pose. The same configuration of eight umbrella sampling was used as described above. For most windows, 20 ns was sufficient since there was minimal interaction between K^+^ and NS11021. However, in the windows centered at 8, 10, 12 and 14 Å, NS11021’s z-distance and orientation with respect to the filter, as well as the K^+^ z-distance to the filter, required extended equilibration over the first ∼15 ns (Figure S7). Consequently, these windows were extended to 100 ns, and the initial 15% of sampling with NS11021 present were discarded prior to analysis.

### Analysis of simulations

The WHAM^77^ was used to calculate the PMFs of K^+^ permeation with the first and second halves of sampling after discarding the initial equilibration region of each window. Each PMF is the mean of these two blocks and the error bars are the difference between the two blocks divided by √2. The bulk PMF was obtained from 200 ns unbiased simulation, with the average of the region > 30 Å set to 0 kcal/mol, and the other PMFs were aligned to the bulk region by a single shift of 0.1 kcal/mol. 2D WHAM was used to reweight and combine all windows and to generate combined PMFs along the biased CV as well as an additional unbiased CV such as pore water count (*N*water) and the orientation of NS11021. The orientation of NS11021 in the pore was characterized using the difference in z-distances from the filter of the CoMs of the FB and AZ groups, ΔZFB-AZ. ΔZFB-AZ < 0 indicates that the FB group is closer to the filter (FB-up), and ΔZFB-AZ > 0 indicates that the AZ group is closer to the filter (AZ-up). When ΔZFB-AZ ≈ 0, NS11021’s long axis is perpendicular to the pore Z axis, and the internal configuration is mainly *syn-syn* (Figure S2). The level of convergence was examined by comparing 2D PMFs derived from the the first and second halves of umbrella sampling trajectories (after discarding the initial 15%). As shown in Figure S9. even though sampling of some of the deep pore orientational states are not fully converged (due to the slow exchange timescale as noted above), the overall PMF is well converged, particularly along the z-distance to the filter dimension (e.g., see the two 1D marginal plots along z-distance).

To estimate the overall *KD* and free energy of NS11021 binding to the pore, we first derived the bound and unbound populations and then calculated *K*D = *Pun*bound/*P*bound and ΔGbind = RT ln *KD*. The z-distance cutoff of 30 Å was used to identify the bound state based on the PMFs (see Figure 4). The errors in KD and ΔGbind, were estimated as the standard error of results calculated from the first and second halves of the umbrella sampling trajectories. To obtain the EC50 of NS11021 at 0 mM Ca^2+^, NS11021 V1/2 data from Rockman, *et al.*^45^ were fit to the log-dose response curve using the Hill equation (Figure S8),

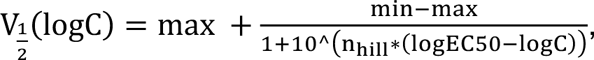

**Figure 3:**
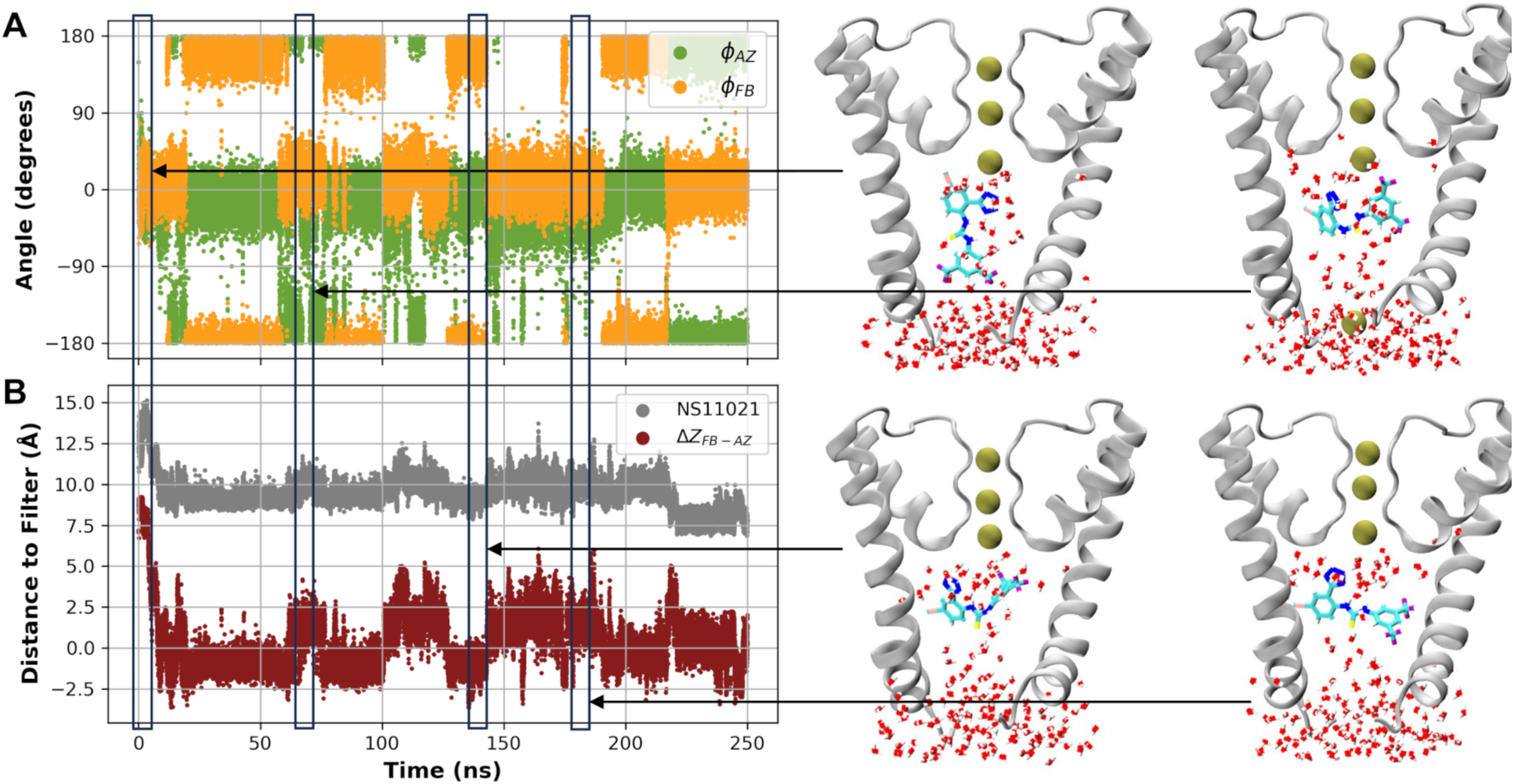
Dynamics of NS11021 in the deactivated BK pore. A) Dihedral angles ΦFB and ΦAZ (see Figure 1B) and **B)** z-distance between the CoMs of NS11021 and the filter, and the orientational CV ΔZFB-AZ, defined by the relative z-distance between the CoMs of the FB and AZ groups to that of the filter. ΔZFB-AZ > 0 indicates AZ-up and ΔZFB-AZ < 0 indicates FB-up.

**Figure 4:**
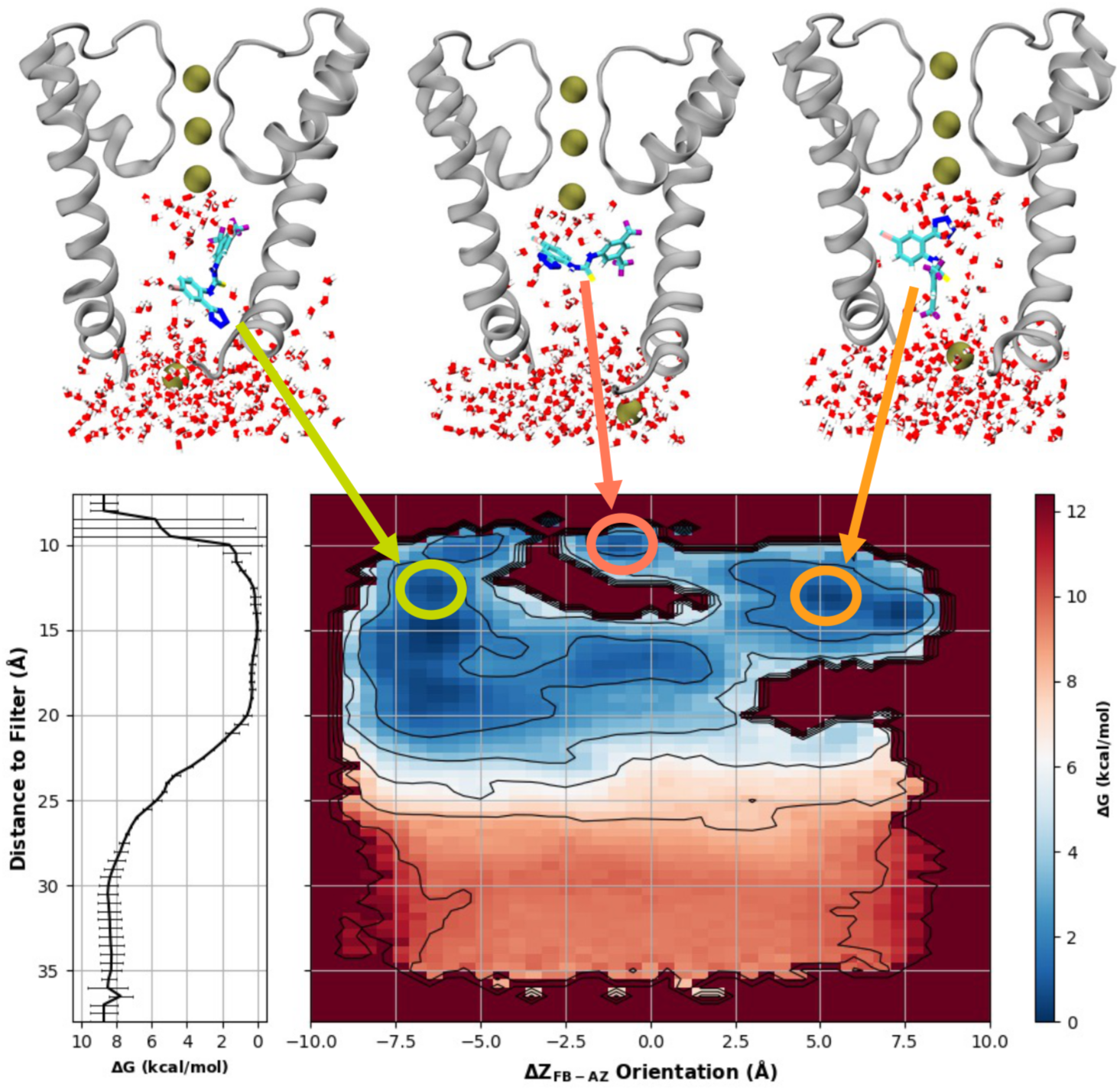
Free energy surface of NS11021 binding to the deactivated BK pore. The 2D PMF was calculated as a function of the orientation of NS11021 (x-axis), described using ΔZFB- AZ (see Figure 3 caption), and the z-distance between the CoMs of NS11021 and the selectivity filter (y-axis). The color bar tick marks and black contours are each distributed in 2 kcal/mol intervals. The marginal plot is the 1D PMF of NS11021 as a function of z-distance from the filter, with error bars showing standard error calculated from block analysis as described in Methods. Representative conformations are shown for representative orientations (FB-up, Az- up and flat) drawn from three minima marked by green, pink and orange circles. The rendering style is the same as in Figure 2.

where *logC* is the log-dose concentration (in μM), *min* and *max* are the minimum and maximum *log*C , and the n_hill_ and *logEC50* are fitted parameters. The fit yielded EC50 = 2.1 μM. All molecular renderings were created with VMD version 1.9.3,^78^ and all data plots were created with matplotlib.^79^

## Results and Discussion

### NS11021 remains dynamic in the pore of BK channels

NS11201 is highly dynamic in solution, with both the AZ and FB groups freely rotatable around the central thiourea group (Figure 1B). A recent solution NMR study of the dihedral conformational propensity of diaryl-thiourea compounds demonstrated that several conformations involving *syn-anti* isomerization are accessible in solution (Figure S2).^56^ Using the dihedral parameters for methylthiourea CHARMM36 force field released in July 2022, we found that indeed NS11021 is dynamic in solution (Figure S3). The least stable state is the most extended *anti-anti* state, which agrees with the NMR data on phenylthiourea compounds^56^ and is consistent with the limited solubility of NS11021.^45^ Equilibrium simulations with NS11021 in various docked conformations revealed that the drug remained highly dynamics inside the deactivated BK pore. As illustrated in Figure 2, not only the internal thiourea dihedrals are quite dynamic in the pore, but also the overall position (z-distance to the filter) and orientation (ΔZFB-AZ) can exchange between various states on the 50 – 100 ns timescale. Figure S10 shows three additional 250-ns equilibrium simulations initiated from different initial configurations, which display similar dynamics in both the internal conformation and the overall binding position and orientation. Such a heterogenous and dynamic bound state of NS11021 in the deactivated BK pore is not surprising considering that the pore is relatively large and hydrophobic and there is a lack of any significant pockets to support specific binding. Note that NS11021 is more dynamic than BL-1249 previously simulated in the pore of TREK-2,^43^ which is a secondary diarylamine with two relatively bulky groups.^80^ Nonetheless, the lack of resolvable density for the bound BL-1249 in the pore suggests that its binding to TREK-2 is likely dynamic as well.

### NS11021 binds the pore via nonspecific hydrophobic interactions

To better characterize the interaction of NS11021, we employed umbrella sampling to sample its conformations and orientations within the confinement of the BK pore. As detailed in Methods, we employed two parallel sets of windows, where the initial configuration of each window was either ΔZFB-AZ > 0 or ΔZFB-AZ < 0. The sampling of all these windows, as well as several windows to sample the bulk solvent region, were combined to produce a single 2D PMF of NS11021 interaction with the BK pore (Figure 4). The PMF highlights a broad conformational landscape with multiple minima of similar free energy levels within the pore, consistent with the observations from the equilibrium MD runs with NS11021 in the pore (Figure 3 and Figure S10). Initial binding is driven primarily by hydrophobic interaction between the hydrophobic FB group and the pore walls, while the charged AZ group remains solvated. As a result, there is a strong preference of the molecule entering the pore in the FB-up orientation. Once NS11021 fully enters the pore (e.g., z-distance < 18 Å), it can sample all three major orientations (FB-up, AZ-up and flat) with similar probabilities. There is only a ‘forbidden region’ in the middle of the pore (z-distance ∼13-14 Å) where ΔZFB-AZ cannot be near 0, due to steric constraints.

It is important to note that the nonspecific and dynamic nature of NS11021 interaction with the pore does not suggest that the binding is weak. The free energy as a function of z-distance to the filter reveals a broad and stable minimum of ∼ -8 kcal/mol from z-distance ∼ 10 – 20 Å (Figure 4). It can be estimated that the total binding free energy ΔGbind = -8.3±0.7 kcal/mol, which is equivalent to *KD* = 0.3-3.1 μM (see Methods). This is in quantitative agreement with an EC50 of 0.4 µM reported by Bentzen, et al.^42^ Using data in Rothberg,^45^ et al. we estimated the EC50 to be 2.1 µM (Figure S8), which again agrees well with the umbrella sampling free energy calculation. We note that directly comparing experimental EC50 range to the calculated *K*D range assumes that the relationship between NS11021 binding and binding-driven activation is approximately linear, which may not be the case. NS11021 may bind to other parts of the BK channel and in more than just the inactivated state. The distribution of open-state dwell times calculated by Rockman, *et al.*^45^ demonstrates that NS11021 could indeed affect both states. However, most of the orthogonal experimental evidence, namely the dramatic change in closed-state dwell times and the increase in the intrinsic *PO*, suggests that NS11021 binds in the closed state. The agreement between our calculated *KD* and experimental EC50 suggests that interaction with the closed pore may dominate the measured effects.

### NS11021 promotes the pore hydrating in the closed state

The pore of BK channels remains physically open with a diameter of ∼ 10 Å in the deactivated state.^39,65,81–84^ It has been proposed that these channels follow the hydrophobic gating mechanism, where the hydrophobic pore undergoes spontaneous dewetting to create a (water) vapor region to block the ion flow.^46–48^ We have previously shown that the free energy cost of hydrating the pore correlates well with the measured gating voltage of mutant BK channels.^48^ We observed that NS11021 binding significantly increase the pore water number from both equilibrium and umbrella sampling simulations (e.g., see Figures 2 and 3), apparently due to the charged AZ group (Figure 1). Using the umbrella sampling simulations of NS11021 binding, we calculated the 2D PMF of NS11021 binding and pore hydration (*N*water) (Figure 5). When NS11021 is outside of the pore, the pore remains stably dehydrated and experienced rare partial hydration events (z-distance > 25). Once NS11021 enters the pore, the hydration free energy of the pore is significantly reduced. There is a new free energy minimum around *N*water ∼ 20 – 30 (and z-distance ∼14 Å), which corresponds to a partially hydrated pore state with NS11021 bound in the inner pore. The increased pore hydration can be expected to shift the pore conformational equilibrium and stabilize the open hydrated pore state as previously suggested by Rockman *et al.*’s electrophysiological experiment and analysis^45^ (also see Introduction).

**Figure 5:**
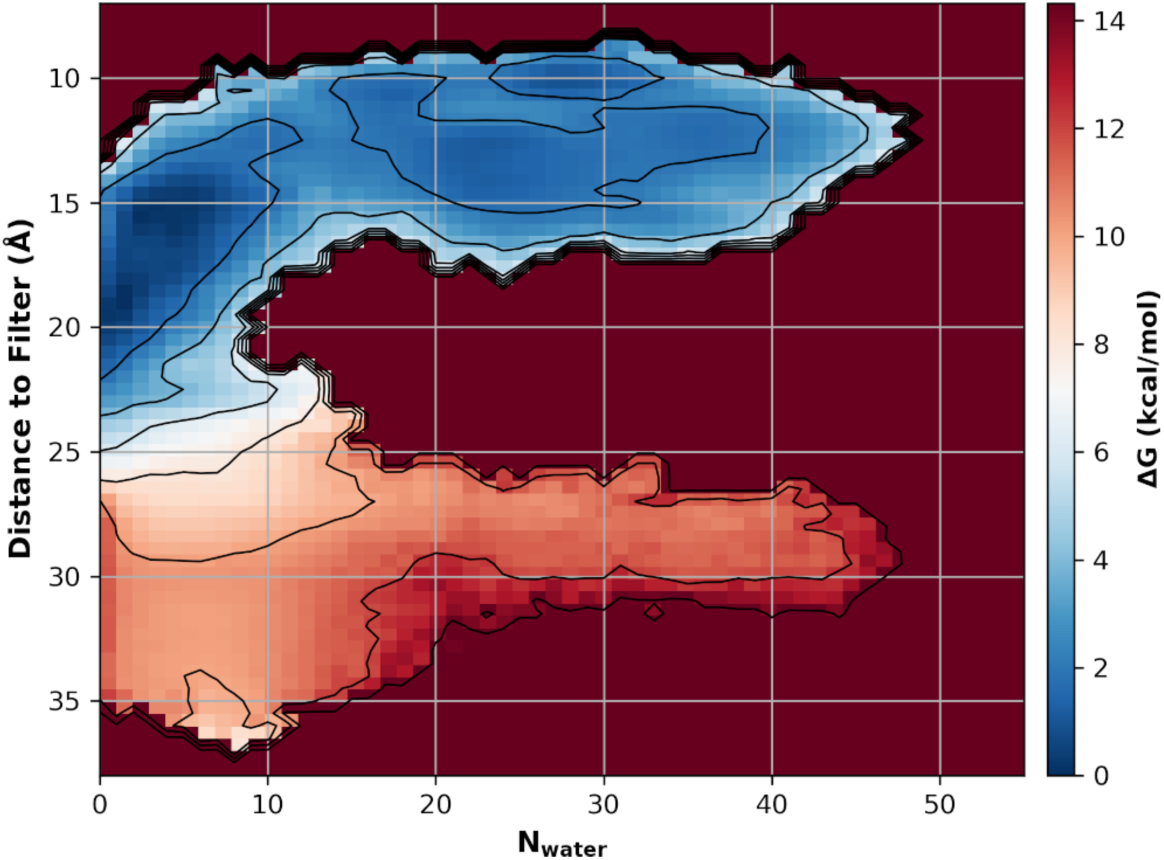
Free energy surface of NS11021 binding and pore hydration. The 2D PMF was calculated as a function of the number of waters in the pore (Nwater) (x-axis) and the z-distance between the CoMs of NS11021 and the selectivity filter (y-axis). The color bar tick marks and black contours are each distributed in 2 kcal/mol intervals.

An important effect of NS11021 on BK channels is the 62-fold increase in intrinsic opening of BK channels.^45^ It has been previously suggested that intrinsic opening is due to the leakage of the vapor barrier in hydrophobic gating,^85^ which can be directly estimated from the free energy barrier of K^+^ permeation through the vapor barrier. For this, we selected three representative free energy basins from the 2D PMF (Figure 4), which represents FB-up, AZ-up and flat orientations, and calculated the PMFs of K^+^ permeation in presence of NS11021 in the deactivated BK pore (see Methods). The results, summarized in Figure 6, show that the barrier is reduced from ∼7.5 kcal/mol to ∼ 3-5 kcal/mol with NS11021 bound. This compares well to the 62-fold reduction in intrinsic opening, which suggests a reduction of ∼ RT ln 62 = 2.5 kcal/mol in the free energy barrier of K^+^ leakage. The modest apparent over-estimation of K^+^ permeation free energy barrier reduction may be attributed to a few factors. The most important is that we did not consider the conformational response of the pore to NS11021 binding, due to the need to restrain the pore structure to achieve good convergence in free energy. It is possible that the pore may contract upon NS11021 binding, making it more difficult for a hydrated K^+^ to leak through. Furthermore, we did not explicitly consider the effect of slowly-exchanging orientations of NS11021 within the pore, which can have nontrivial effects on the K^+^ permeation through the deactivated pore.

**Figure 6:**
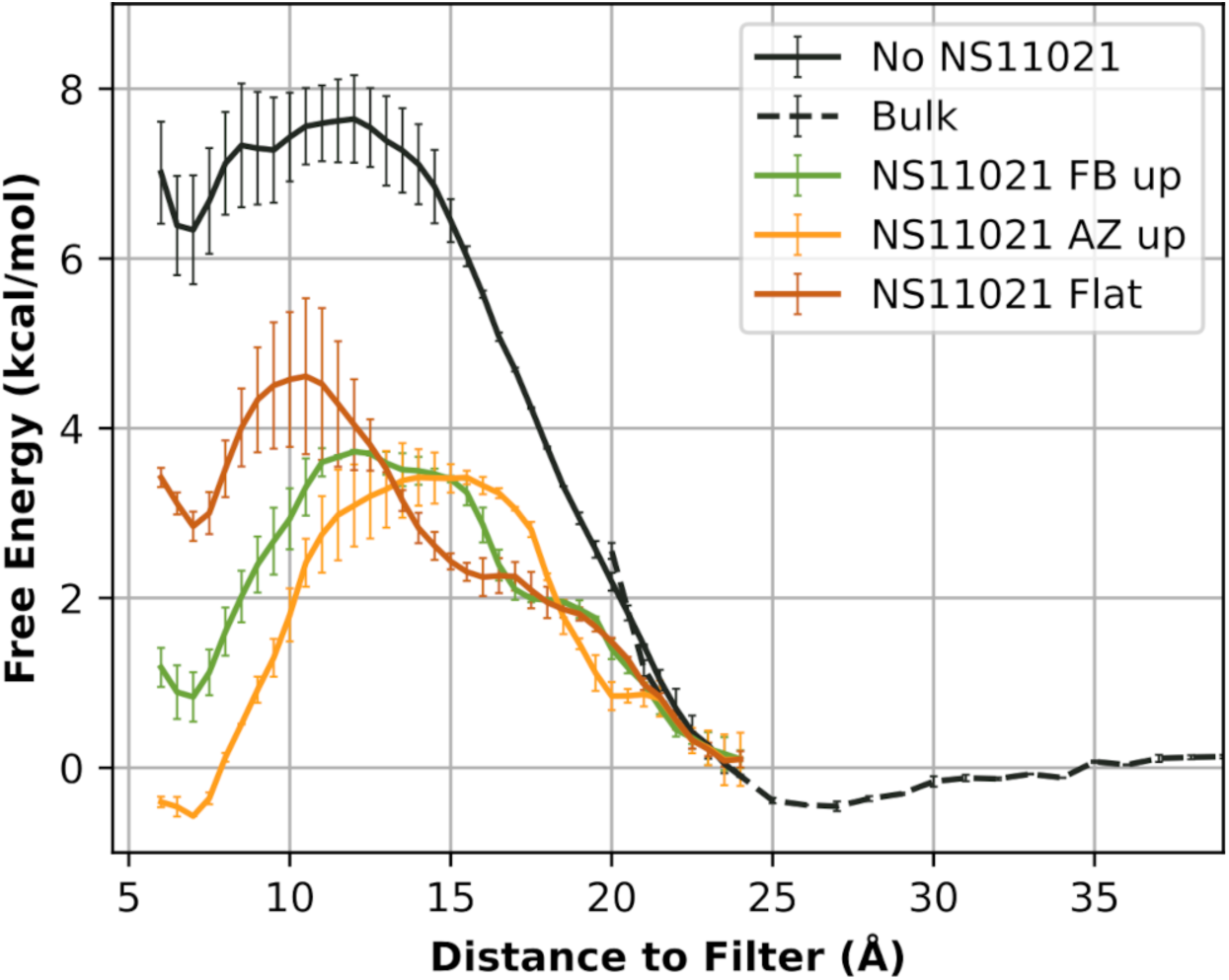
Free energy of K^+^ permeation with NS11021 bound in the pore in three representative configurations. NS11021 restrained to the representative states shown in Figure 4 (FB-up, AZ-up and flat). The black curve is the PMF without NS11021 in the pore as a reference, and the dashed curve is the PMF extending away from the protein into the bulk solute derived from unbiased simulations.

## Conclusions

Atomistic simulations and free energy analysis were performed to examine the molecular detail of binding and possible activation mechanism of a small molecule activator NS11021 on the BK channel. The results show that the molecules initially contacts the pore with the hydrophobic FB end facing into the dewetted, hydrophobic pore in the deactivated state. Once inside the inner pore, NS11021 remains dynamic and sample a wide range of conformational and orientational states. The dynamic binding mode is consistent with a lack of resolvable electron density in the X-ray crystallography despite mutational data supporting the pore binding. Despite the dynamic nature, the drug molecule makes extensive hydrophobic contacts with the inner pore. The effective KD of 0.3-3.1 μM derived from the free energy analysis agrees well with the experimental EC50 range of 0.4-2.1 µM.^42,45^ The most prominent effect of NS11021 binding seems to be a substantial increase in the pore hydration. BK channels have been suggested to follow the hydrophobic gating mechanism, where the pore does not physically close but rely on hydrophobic dewetting to create a vapor barrier to block ion permeation.^46–48^ As such, increased pore hydration by NS11021 likely drives the open-close equilibrium of the channel towards the open state suggested by electrophysiological analysis. It has been previously shown that intrinsic opening of BK channels likely derive from leakage of vapor barrier in hydrophobic gating.^85^ Therefore, increased pore hydration level in the presence of NS11021 binding likely also explains the large 62-fold increase in intrinsic opening of the BK channel. Indeed, K^+^ permeation PMFs calculated with NS11021 restrained in three distinct and representative orientations reveal significant decrease of K^+^ leakage barrier, by 3-5 kcal/mol. Taken together, the current study provides new insights on the molecular basis of the activation effect of NS11021 on BK channels. The proposed mechanism of promoting hydration is largely insensitive to the NS11021 binding pose within the pore. Instead, it only requires nonspecific hydrophobic interactions mediated by the nonpolar moieties the drug molecule to bring the negative charged group into the pore and increase hydration propensity. Such a general mechanism could explain why the class of NCAs is able to activate many different K^+^ channels. This knowledge will inform the use of NCAs in studies of K^+^ channel function and may also prove useful in future efforts to directly target the hydrophobic gate via therapeutic agents.

## Acknowledgements

The simulations in this study were run on the pikes cluster housed in the Massachusetts Green High-Performance Computing Cluster. This work was supported by National Institutes of Health Grant R35 GM144045 (J.C.). E.B.N. was also supported by National Research Service Award T32 GM139789 from the National Institutes of Health.

## Author contributions

All authors contributed to conceiving of the research and writing and revising the manuscript. E.B.N and Z.J. performed the simulations and analyses.

## Supporting Information

Illustrations of NS11021 charge parameters, naming conventions, and dihedral free energy surface in bulk and in BK pore; traces and histograms of NS11021 and K^+^ permeation umbrella sampling simulations; log dose-response curve of NS11021 using data from reference^45^; convergence of 2D free energy surface of NS11021 orientation; and traces of unbiased equilibrium simulations of NS11021 in the BK pore.

---

**Figure S1:**
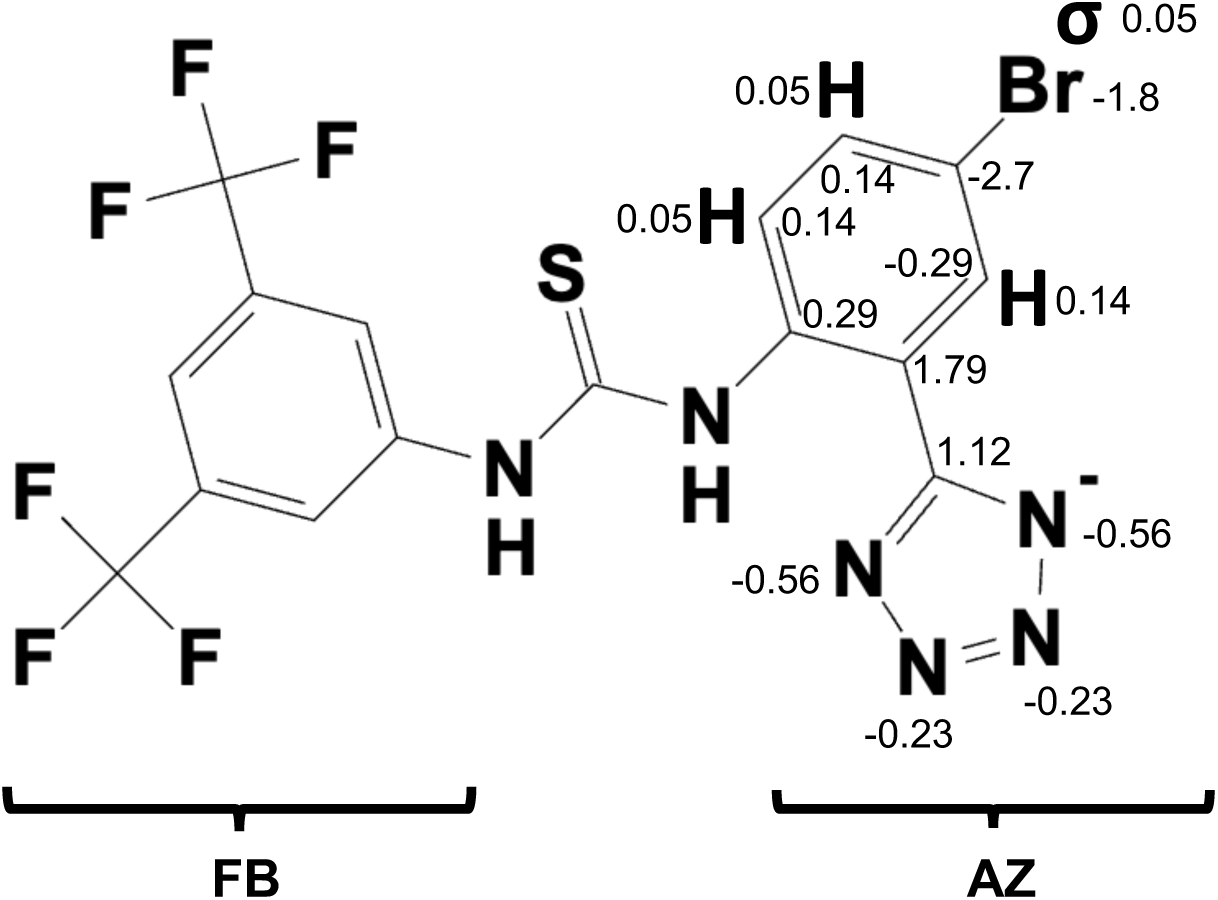
Illustration of charges on tetrazol-phenyl group. The charges for the tetrazol- phenyl (AZ) group were determined following the standard CGenFF protocol, as described in the main text. The AZ and di-trifluoromethyl-phenyl (FB) groups are denoted with red text and brackets.

**Figure S2:**
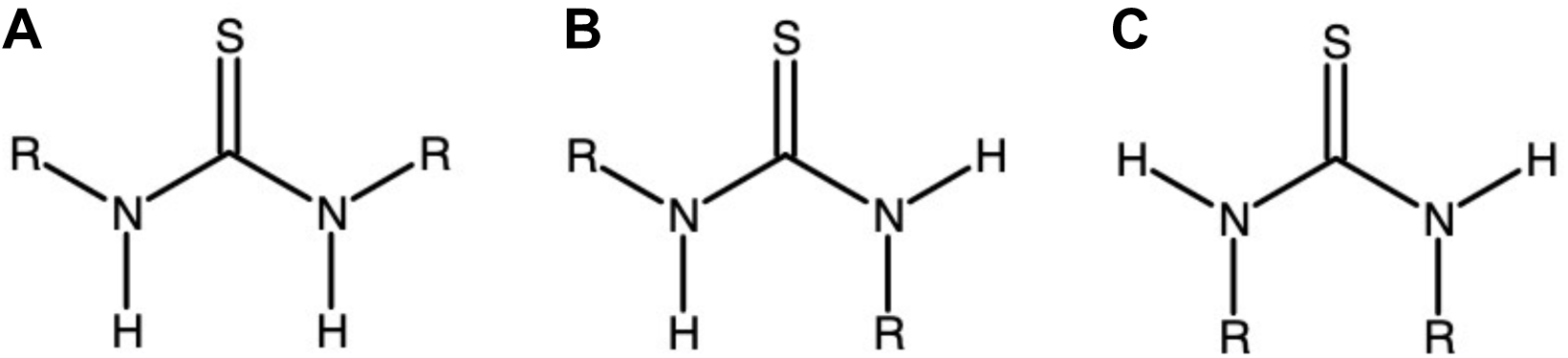
Illustration of diaryl-thiourea torsion nomenclature. These diagrams depict **A)** *anti-anti*, **B)** *anti-syn*, **and C)** *syn-syn*, respectively.

**Figure S3:**
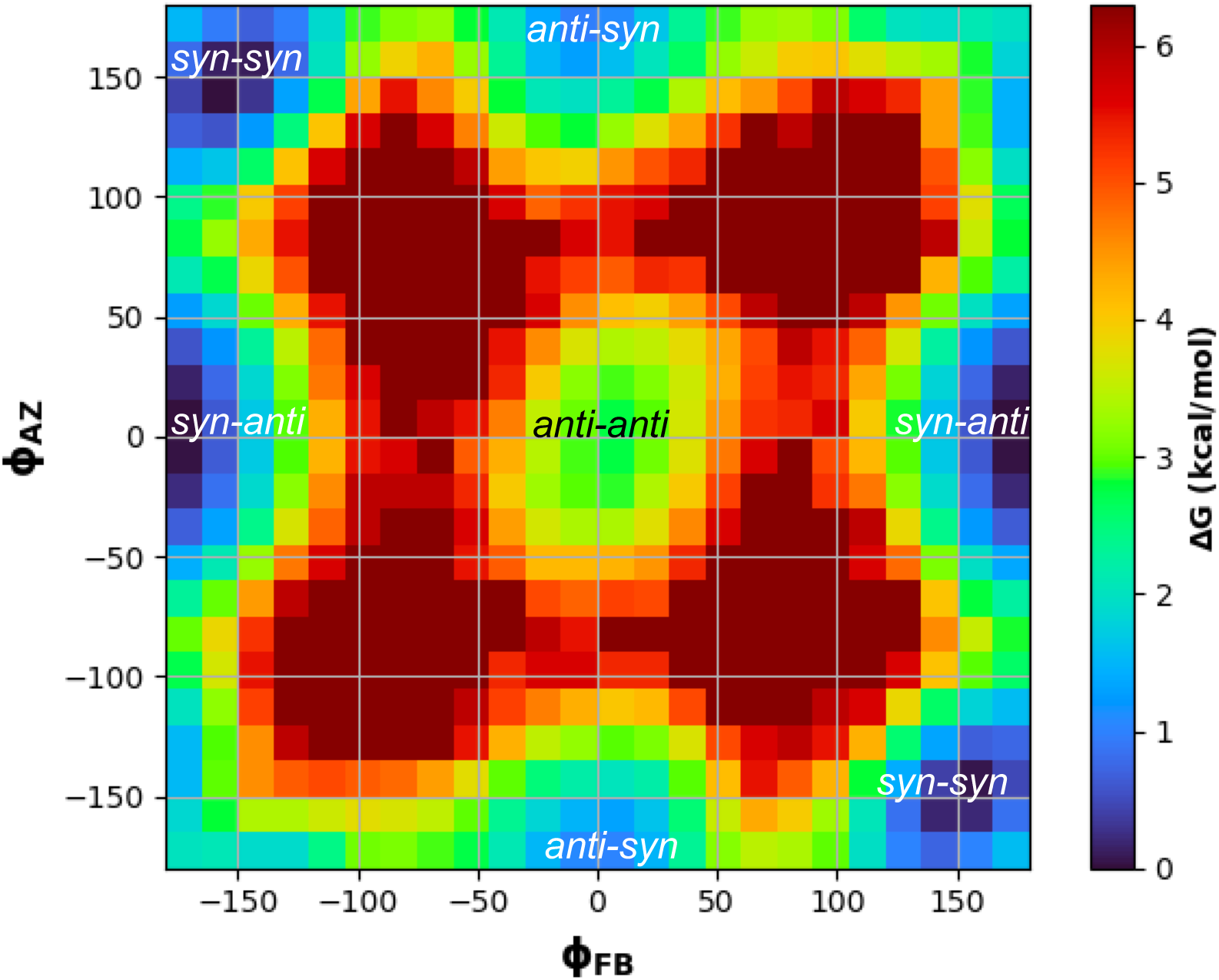
NS11021 dihedral free energy surface. Derived from 2D histograms calculated from a 100 ns simulation of a single NS11021 in TIP3 water. (Φ_!"_, Φ_#$_) = (0,0) corresponds to the *anti-anti* configuration depicted in Figure S2.

**Figure S4:**
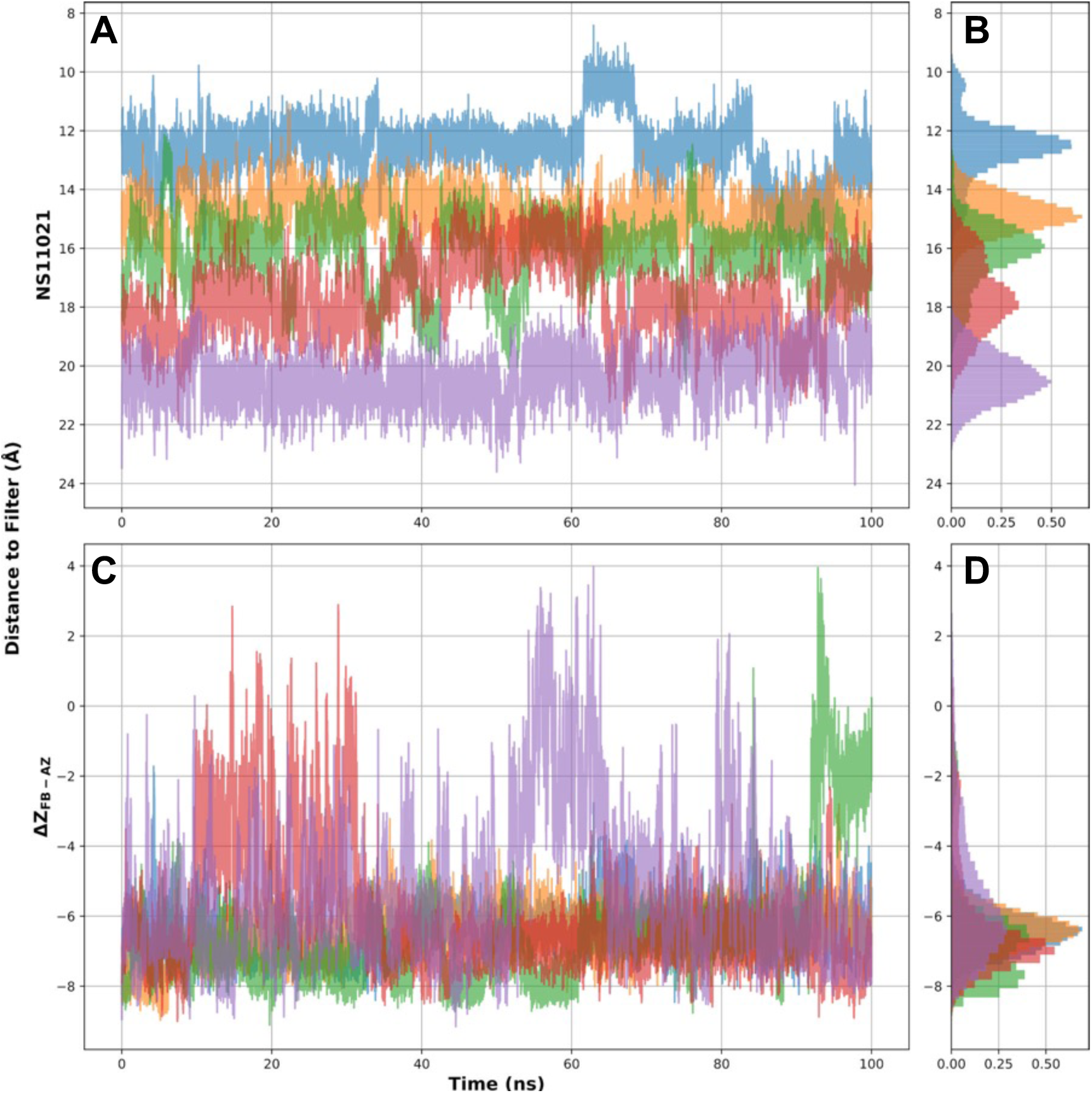
Time traces and histograms from umbrella sampling of NS11021 with a FB- up initial orientation. Panels A and B show the trace and histogram of the CoM distance between the NS11021 and the filter. Panels C and D show the trace and histogram of the NS11021 orientation CV (ΔZFB-AZ).

**Figure S5:**
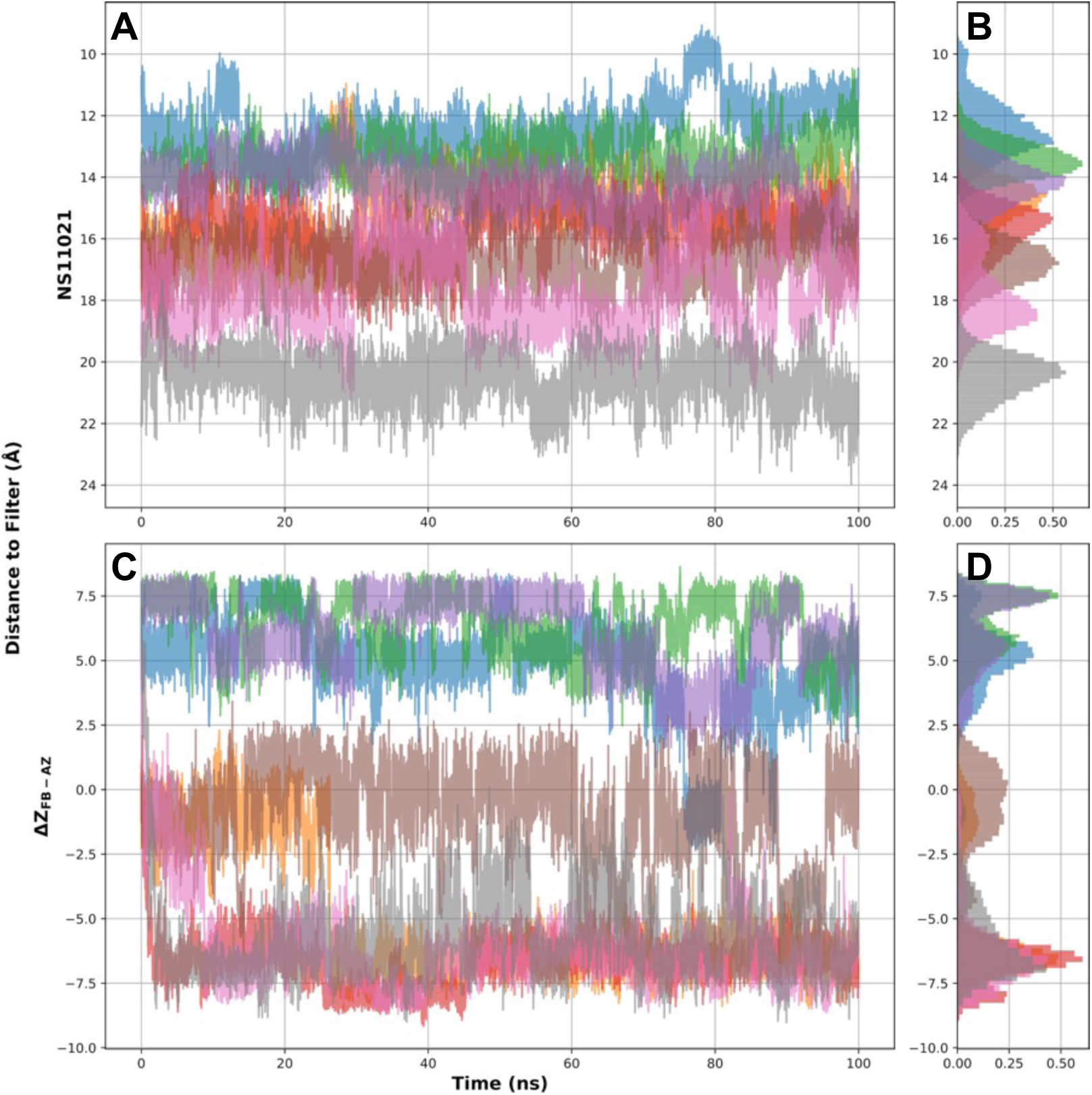
Time traces and histograms umbrella sampling of NS11021 with an AZ-up initial orientation. Panels A and B show the trace and histogram of the CoM distance between the NS11021 and the filter. Panels C and D show the trace and histogram of the NS11021 orientation CV (ΔZFB-AZ).

**Figure S6:**
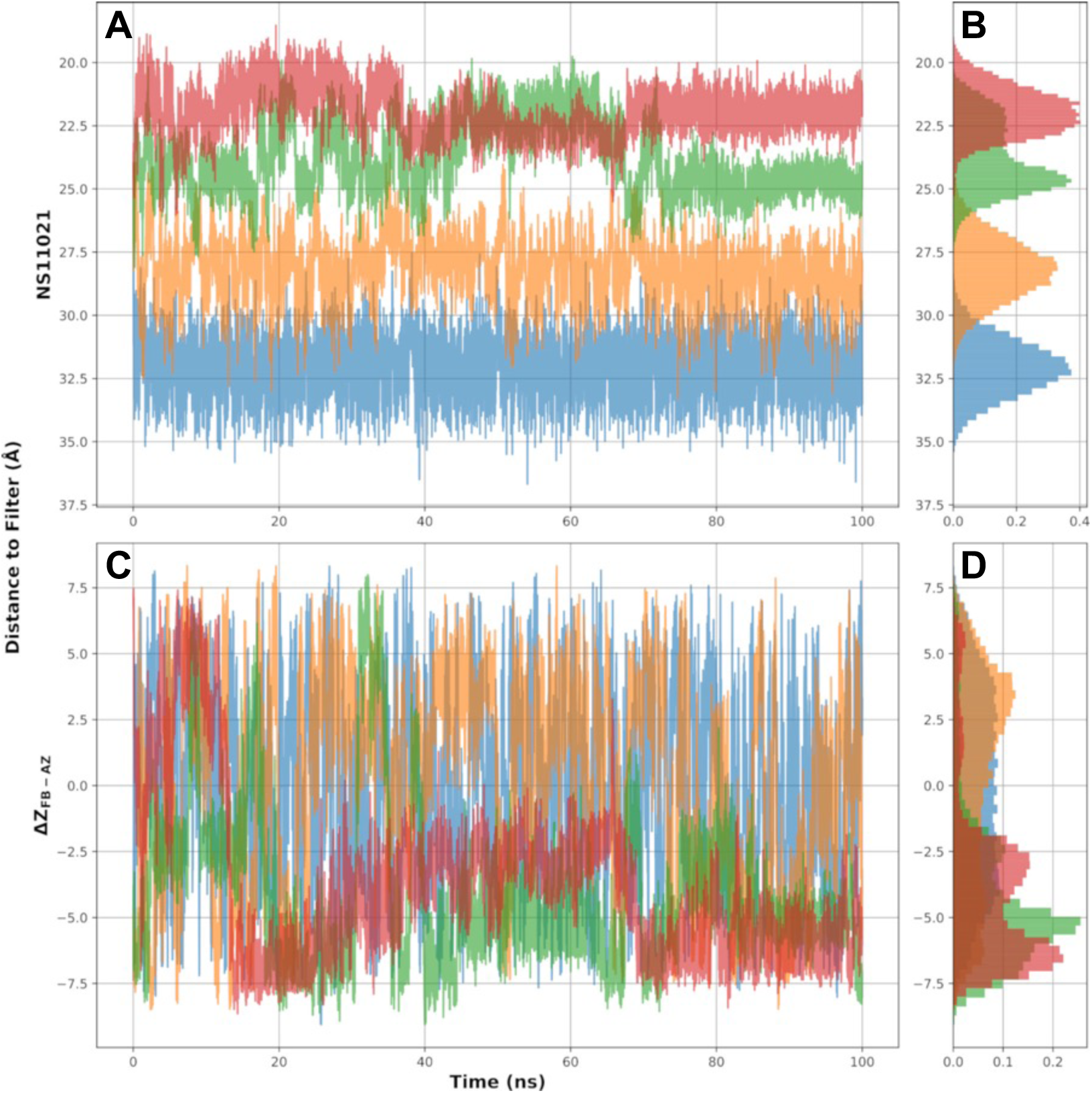
Time traces and histograms umbrella sampling of NS11021 outside the pore. The initial orientation of NS11021 in these windows were AZ-up. Panels A and B show the trace and histogram of the CoM distance between the NS11021 and the filter. Panels C and D show the trace and histogram of the NS11021 orientation CV (ΔZFB-AZ).

**Figure S7:**
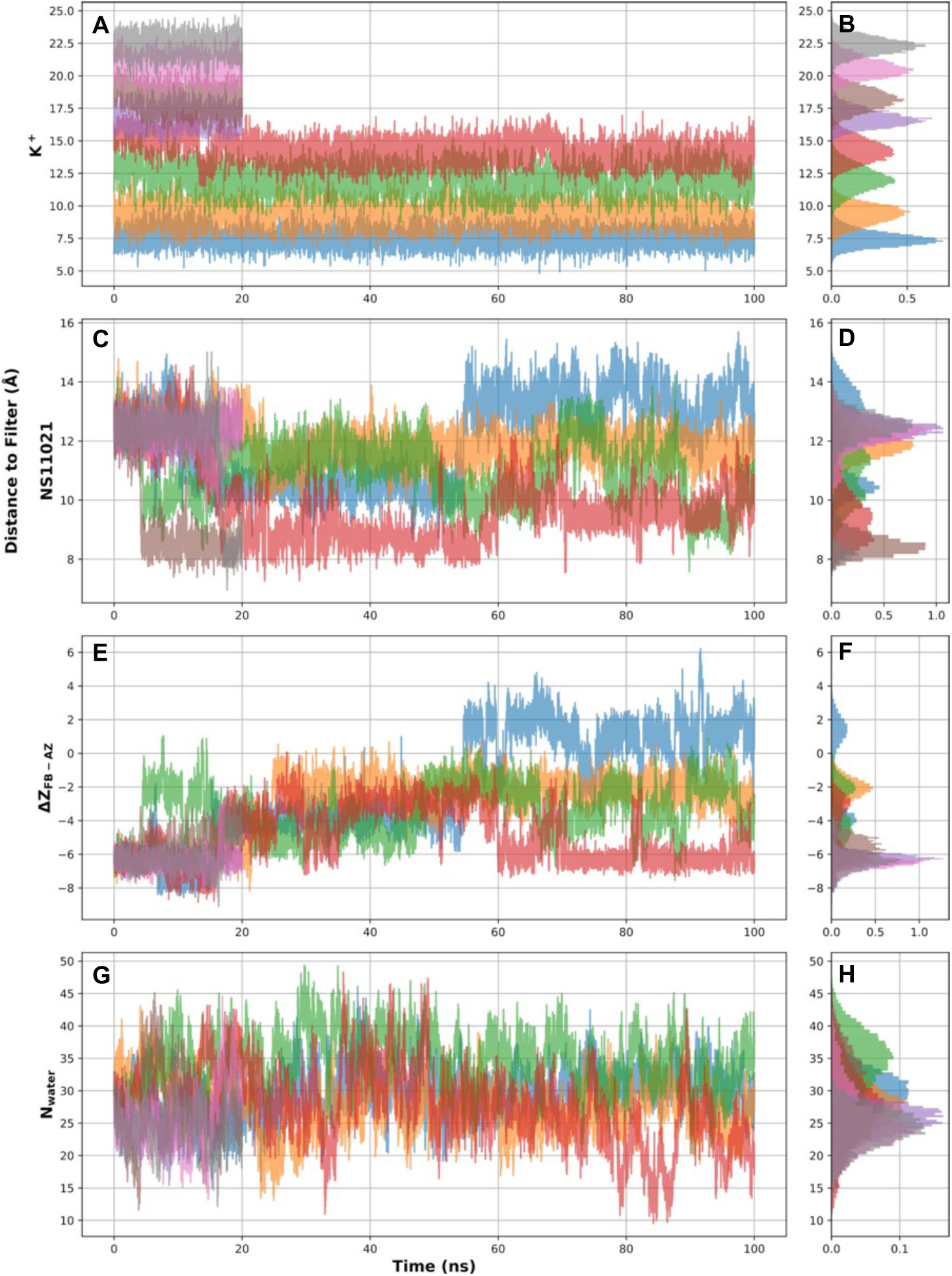
Time traces and histograms from umbrella sampling of K^+^ permeation with NS11021 in the pore. These data come from windows where NS11021 was harmonically restrained to its initial CoM. The initial orientation is FB-up. Panels A, C, and E show traces of the K^+^, NS11021 CoM, and NS11021 orientation (ΔZFB-AZ) with respect to the filter, and panels B, D, and F show the corresponding histograms to panels A, C, and E, respectively. Panel G shows the trace of water in the pore, and panel H shows the corresponding histogram.

**Figure S8:**
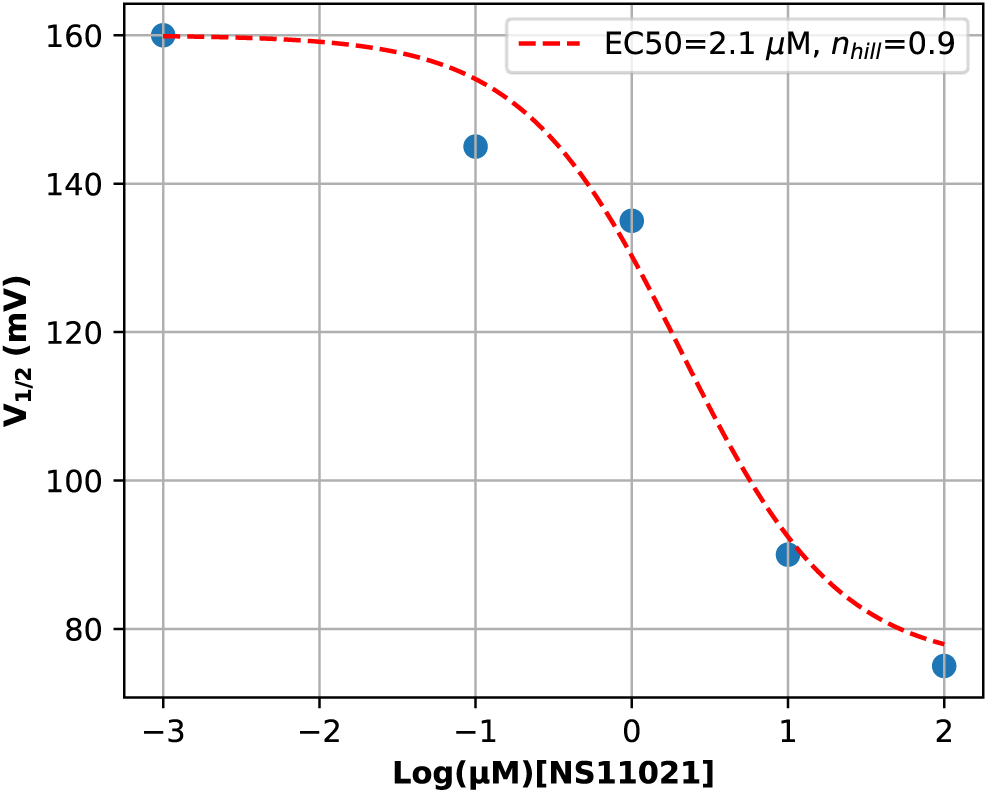
Log dose-response curve for EC50 estimation. The data (blue dots) was obtained from Rockman, *et al.* (2020) (0 mM [Ca^2+^]) and fit with the Hill equation (red line). The lowest concentration was nominally 0 µM NS11021, but to facilitate readability we plotted it as nM or 10^-3^ µM.

**Figure S9:**
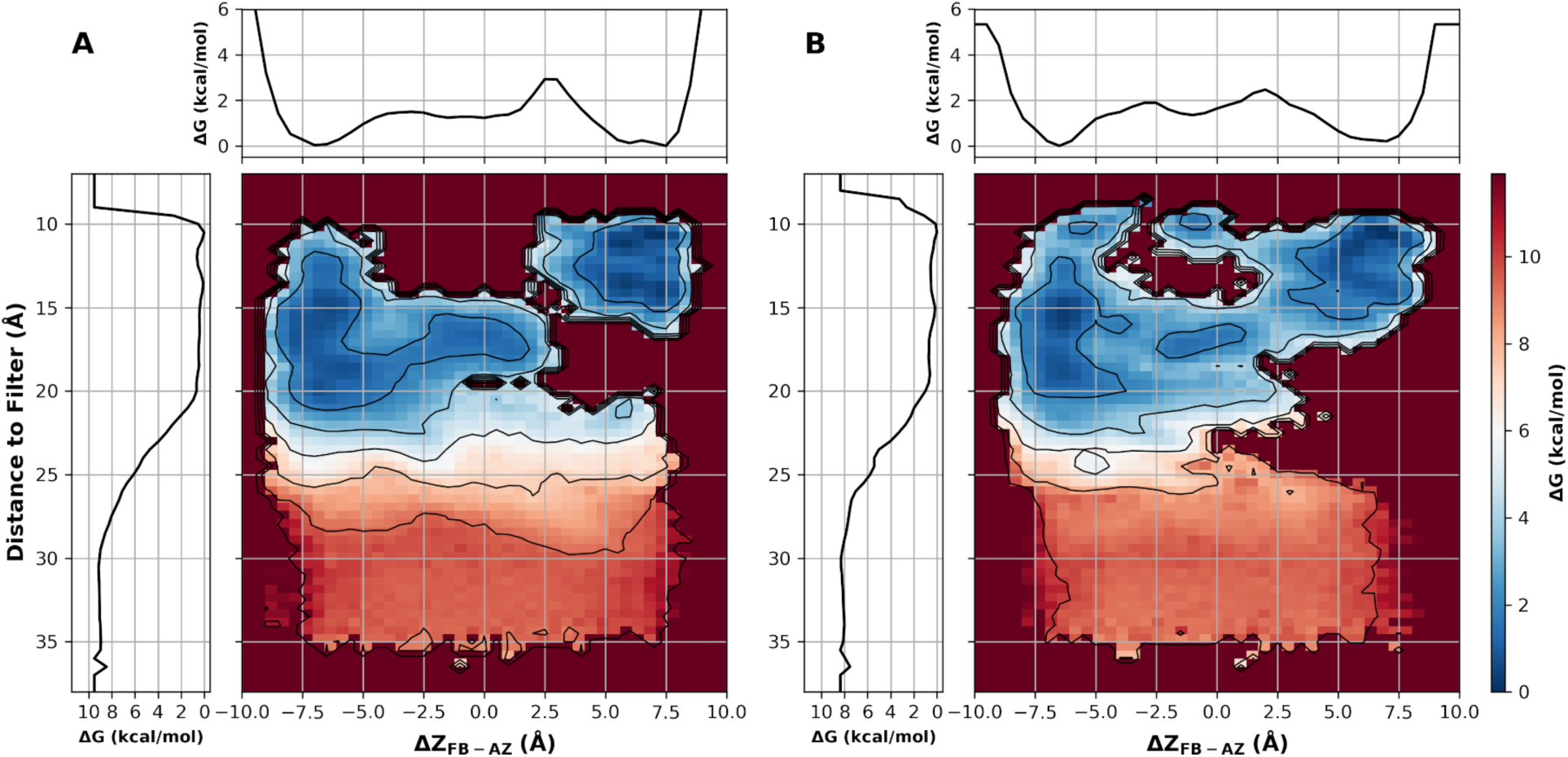
Convergence of the free energy surface of NS11021 binding to the deactivated BK pore. Panel A comes from the first half of the windows and panel B comes from the second half, after discarding the initial 15 ns of each window.

**Figure S10:**
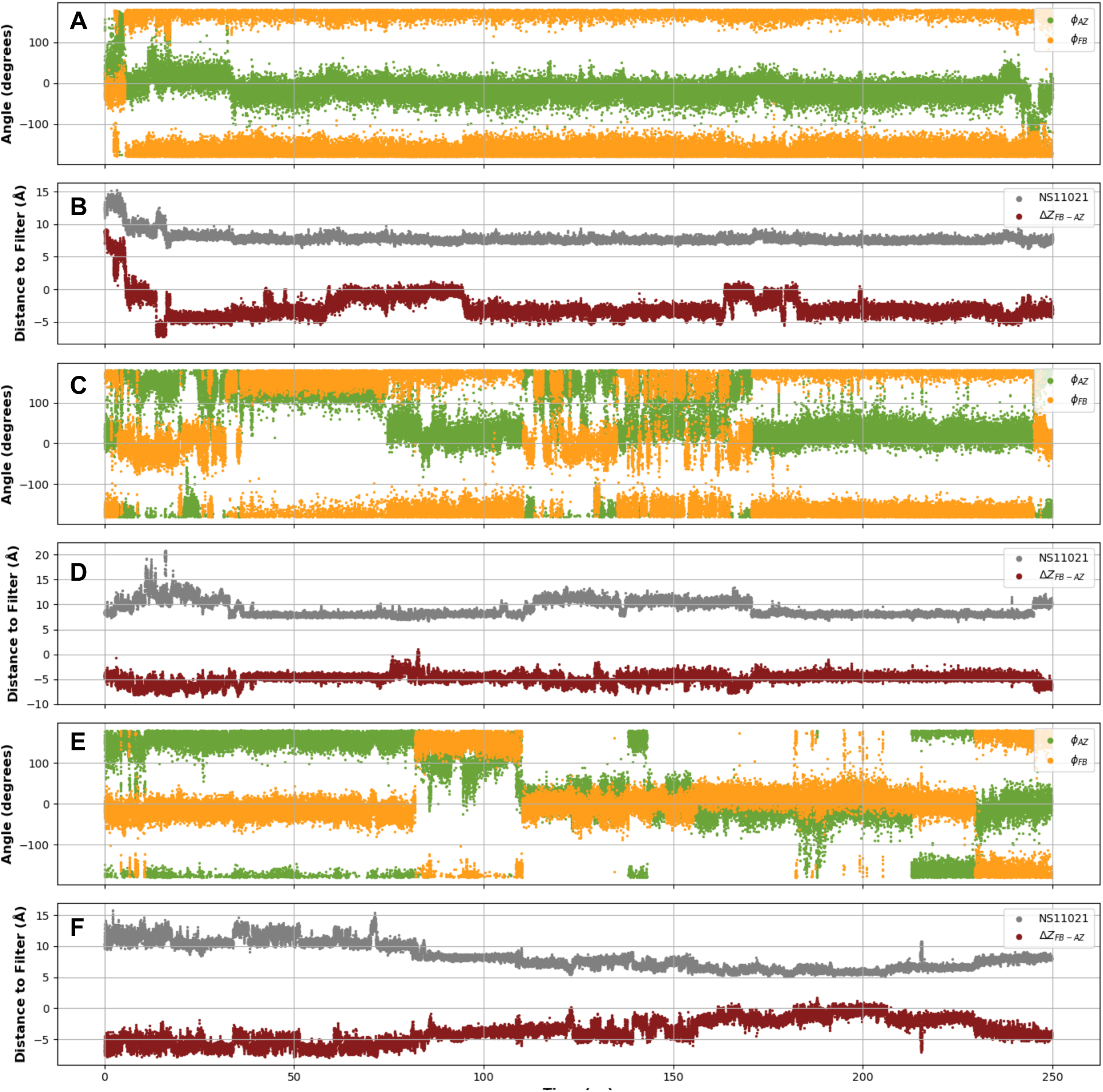
Traces of three independent simulations of NS11021 in the pore. Panels A, C, and E show the internal thiourea dihedral angles Φ_!"_ and Φ_#$_, and panels B, D, and F show the z-distance between the CoM of NS11021 and the filter, as well as the orientation CV ΔZFB- AZ. The simulation in panels A and B was initiated with an AZ-up orientation, and simulations in panels C-F were initiated with FB-up orientations.

